# Emergence of distinct neural subspaces in motor cortical dynamics during volitional adjustments of ongoing locomotion

**DOI:** 10.1101/2022.04.03.486001

**Authors:** David Xing, Wilson Truccolo, David A. Borton

## Abstract

3

The brain is capable of simultaneously carrying out multiple functions, such as making different types of movements at the same time. One example is how we are able to both carry out stereotyped walking or running movements, while concurrently performing precise, target-directed movements such as kicking a ball in a soccer match. Recently, research has shown that different computations within the same population of neurons can be carried out without disrupting each other by confining the processes into separate subspaces. Whether this strategy is used to precisely control our limbs while maintaining locomotion is still an open question. Here, we recorded the activity of primary motor cortex in nonhuman primates during obstacle avoidance on a treadmill. We found that the same neural population was active during both basic unobstructed locomotion and volitional obstacle avoidance movements. Additionally, we identified the neural modes spanning the subspace of the low-dimensional dynamics in M1 using both supervised and unsupervised techniques. We found that motor cortex employs a subspace that consistently maintains the same cyclic activity throughout obstacle stepping, despite large changes in the movement itself. All the variance corresponding to the large change in movement during the obstacle avoidance is confined to its own distinct subspace. Our findings suggest that M1 utilizes different activity subspaces to coordinate the maintenance of ongoing locomotor-related neural dynamics and fast volitional gait adjustments during complex locomotion.

**Significance Statement:** Our ability to modulate our ongoing walking gait with precise, voluntary adjustments is what allows us to navigate complex terrains. Locomotion and precise, goal-directed movements, such as reaching are two distinct movement modalities and have been shown to have differing requirements of motor cortical input. It is unknown how these two movements are represented in M1 low dimensional dynamics when both are carried out at the same time, such as during obstacle avoidance. We developed a novel obstacle avoidance paradigm in freely-moving non-human primates and discovered that the strategy employed by motor cortex is to confine the rhythmic locomotion-related dynamics and the voluntary, gait-adjustment movement into separate subspaces.

## 5 Introduction

The nervous system is a highly flexible computational machine that is capable of simultaneously performing many different functions. An emergent view on neural computation posits that networks of neurons engage in specific patterns of covariation for carrying out specific functions (Athalye et al., 2017; Churchland et al., 2012; Elsayed et al., 2016; Gallego et al., 2018; Kaufman et al., 2014; Mante et al., 2013; Pandarinath et al., 2018; Stavisky et al., 2017). By constraining these coactivation patterns, which we will refer to as *neural modes*, to separate subspaces, the network is able to carry out computations associated with one function without interfering with the activity related to different functions in the other subspaces. This allows for the same neural population to carry out multiple different processes. For example, in order to avoid prematurely activating downstream muscles during movement preparation, the neural variance in motor cortex resides primarily within an output-null neural subspace during the quiescent preparatory period, before transitioning into the distinct, output-potent subspace once the muscles becomes active during movement(Kaufman et al., 2014). By segregating the neural activity into distinct subspaces, the same network is able to engage in both movement preparation and movement execution.

It is still unknown whether within movement execution itself, different types of movement performed simultaneously may also correspond to distinct subspaces. One particular behavior that may be amenable to the subspace-segregation hypothesis is locomotion. Previous studies have shown that the oscillator circuits responsible for basic locomotor movements reside in the spinal cord (McCrea & Rybak, 2008), and descending input from higher cortical areas such as motor cortex is not required for carrying out unobstructed walking on a treadmill (G raham Brown, 1911; Grillner et al., 1997). However, many animals have also developed the ability to precisely position their limbs in specific locations and orientations during locomotion, which facilitates navigation in complex terrains (Beloozerova & Sirota, 1998; Georgopoulos & Grillner, 1989; Porter & Lemon, 1995; Yakovenko & Drew, 2015). During these behaviors, information about the environment is used to generate specific volitional movements which are integrated with the underlying locomotion rhythm. These volitional, gait-modifying movements are distinct from basic unobstructed locomotion which consists of repeated rhythmic movements agnostic to the environment. Unlike in basic locomotion, motor cortex is essential for carrying out precise adjustments to the movements of the limb (Beloozerova & Sirota, 1998; Courtine, 2005; Drew et al., 2002; Liddell & Phillips, 1944) and must integrate top down control of the muscles with incoming afferent information from downstream areas (figure 1a) (Petersen et al., 1998; Pruszynski et al., 2011; Scott et al., 2015). How M1 carries out the necessary computations to produce the correct volitional movement while also accounting for the underlying locomotion movements is poorly understood, especially in primates. One possibility could be that motor cortex engages separate neural subspaces for tracking the basic cyclic rhythm and for carrying out the targeted, goal-oriented movements (figure 2b). This mechanism allows the same neural circuit to perform targeted, visually-guided movements, such as those studied in classical center-out reaching paradigms, while at the same time preserving underlying walking movements during complex tasks such as walking across stepping stones or performing obstacle avoidance.

**Figure 1:**
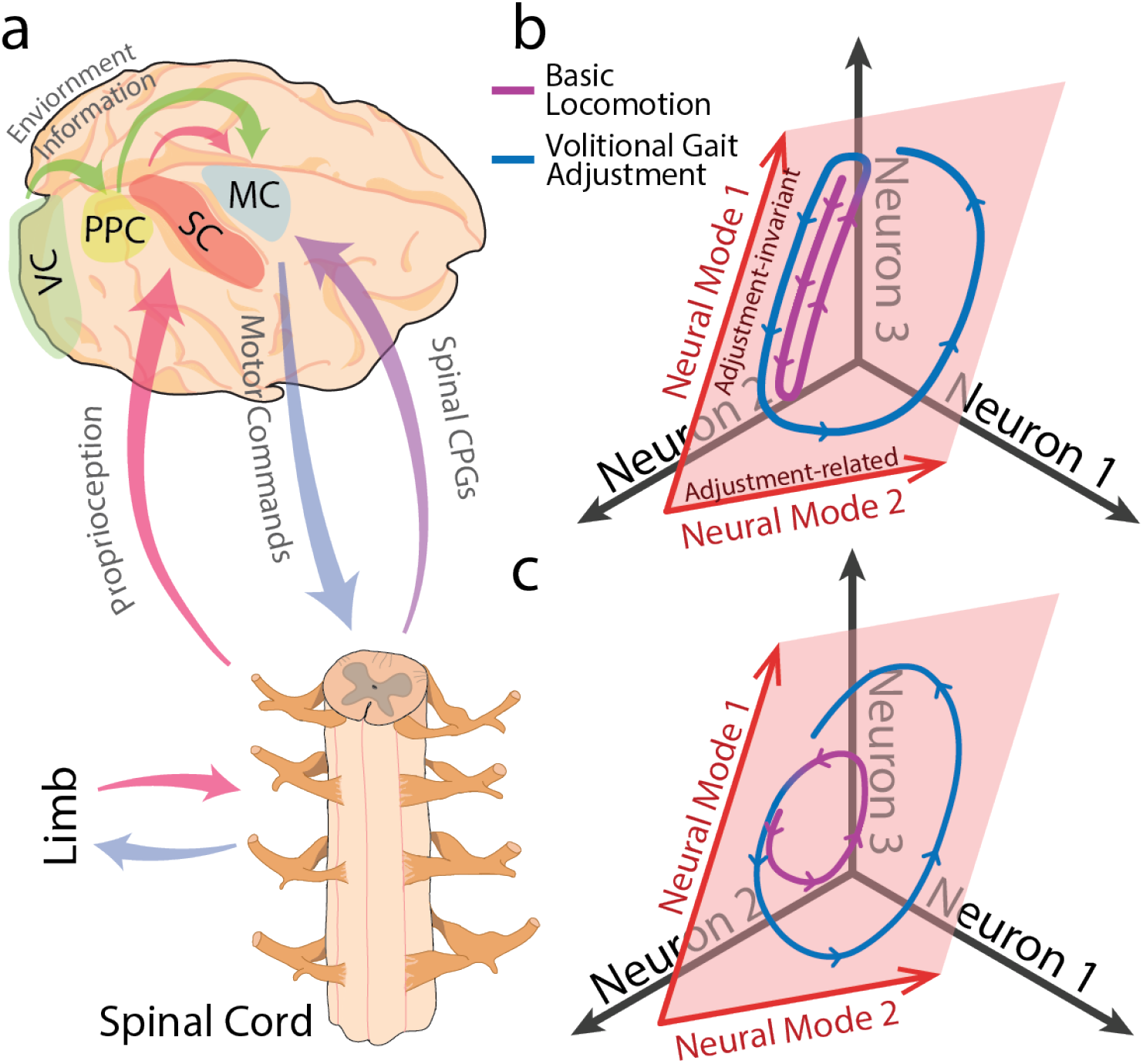
Neural modes related to volitional adjustments during locomotion. **a)** Example pathways of some of the converging inputs onto motor cortex during volitional gait adjustments. Proprioception and spinal CPGs provide information about the limb position in the gait cycle while visual information from the environment both drives decision making to react to complex environments as well as facilitating the calculation of the precise movement needed to successfully navigate the terrain. The nervous system must integrate all of these components to generate the appropriate motor command, which is relayed through the spinal cord to the limb. MC - motor cortex, SC - somatosensory cortex, VC - visual cortex, PPC - posterior parietal cortex. **b**,**c)** Two possible strategies that motor cortex could employ to carry out volitional gait adjustments. The full three dimensional space represents all possible combinations of firing rates of three toy neurons. The neural activity is confined to a two-dimensional subspace (red plane) spanned by two neural modes (red arrows). The curve represents the time-varying neural activity during one stride of basic locomotion (purple) followed by a stride with a volitional gait adjustment (blue). In **b**, the neural activity during basic locomotion is mostly confined to the first neural mode, while the second neural mode encodes the movement modifications. In **c**, both neural modes are utilized during basic locomotion and both are modified by motor cortex during the volitional movement.

**Figure 2:**
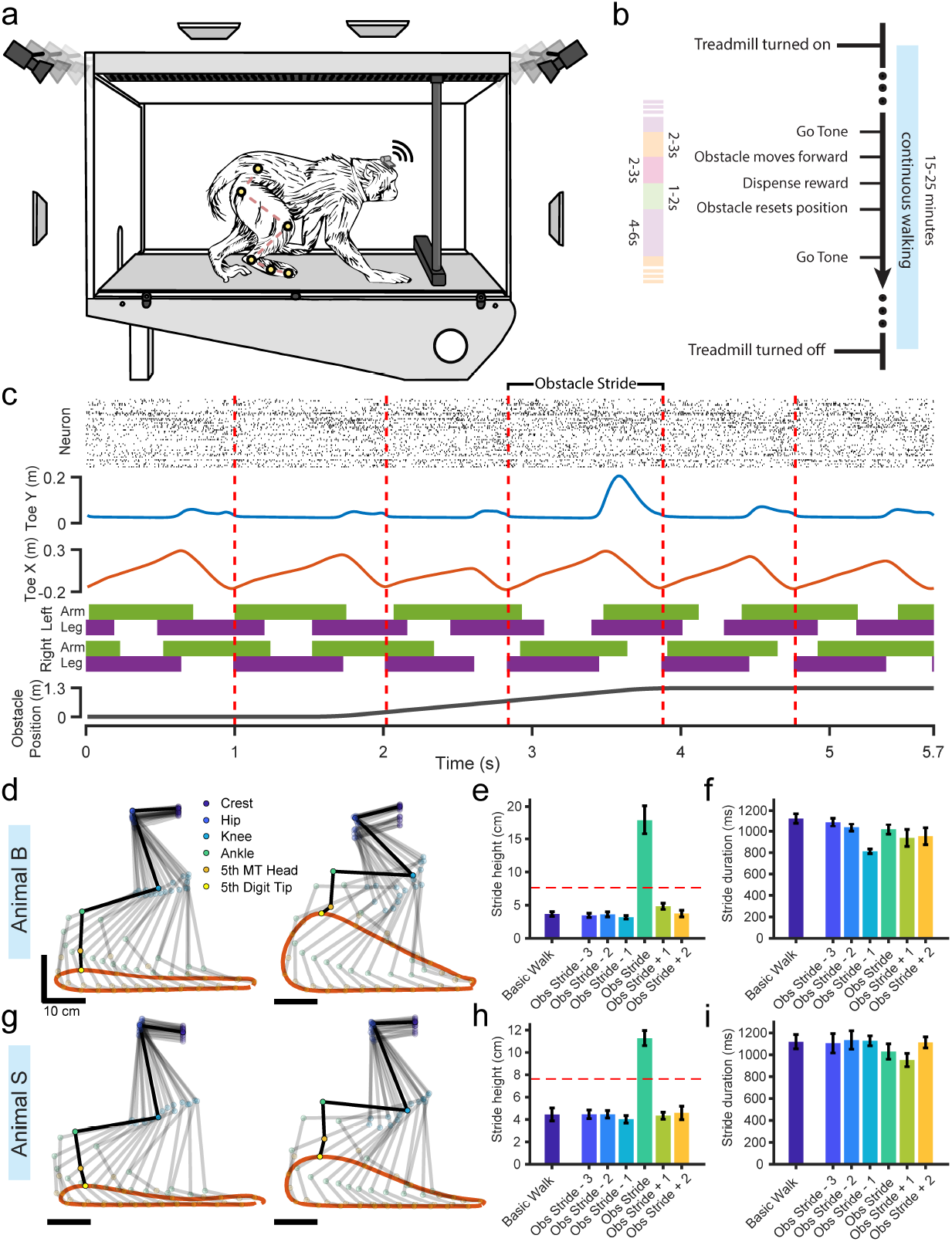
Experimental paradigm and behavior. **a)** Treadmill enclosure and behavioral apparatus where animals performed locomotion and obstacle stepping. Antennas surrounding the enclosure wirelessly collected neural data while video cameras recorded the positions of six joint markers painted on the hind limb. **b)** Behavioral paradigm. Each trial consisted of a single obstacle run. **c)** Example data from a single trial of animal B, showing the stride bringing the leg over the obstacle along with three strides before the obstacle stride and two strides after the obstacle stride. Top: raster plot of sorted single units from the implanted Leg-M1 MEA. Blue trace: height of the right toe tip. Orange trace: horizontal position of the right toe tip, normalized to the right illiac crest. Second to bottom: gait pattern across all four limbs. Solid bars represents contact with the ground (stance phase); purple - hindlimbs, green - forelimbs. Bottom: Obstacle position along the path of the treadmill. **d**,**g)** Stick diagram of the right hind limb during one stride of basic, unobstructed walking (left) or during the stride stepping over the obstacle (right). Orange trace represents the trajectory of the toe tip. Stick figures are spaced 60ms apart, with the dark stick figure highlighting the limb during the maximum height of the toe tip. **e**,**h)** Maximum height reached by the toe tip for either continuous unobstructed (basic) walking or the stride over the obstacle and the surrounding strides. Error bars represent standard deviation. Dotted red line indicates the height of the obstacle. **f**,**i)** Duration of each stride, error bars are standard deviation. **d-f** : animal B, **g-i**: animal S.

Here, we recorded neural activity from the leg area of M1 in freely-moving non-human primates while they performed basic locomotion as well as stepping over an incoming obstacle on a treadmill. We found that locomotion movements, as well as volitional, gait-adjusting movement are represented in the same recorded neural population. Using dimensionality reduction, we found underlying neural modes which are completely unaffected by the obstacle avoidance movement, despite large changes in the kinematics. Separate neural modes captured the variance in the population activity encoding the precise obstacle avoidance movement. Therefore, M1 appears to engage two distinct subspaces, one for maintaining the ongoing cyclic dynamics present during locomotion, and one for encoding the momentary change in M1 engagement during the gait modification.

## 6 Methods

### 6.1 Animal husbandry

All experimental and surgical procedures were performed under approved Institutional Animal Care and Use Committee (IACUC) protocols at Brown University. Two male rhesus macaques, aged 7 and 9 years old were housed in individual cages and trained to perform the obstacle avoidance walking task. Positive reinforcement in the form of solid food (Starbursts candy, yogurt covered raisins), was used and primate biscuits and water were provided *ad lib* throughout training.

### 6.2 Experimental design

All behavioral tasks were carried out inside a treadmill enclosure. The treadmill was purchased commercially (JogADog, MI), and a custom plexiglass box (177.8cm long x 47.6cm wide x 91.4cm tall) was constructed above it (figure 2a). Similar to previous recording studies (e.g. (Berger et al., 2020; Foster et al., 2014; Yin et al., 2014)), animals were able to move freely inside the enclosure and were not tethered in any way. To encourage consistency of movements across trials and also to protect the obstacle components of the apparatus, plexiglass walls were placed below the ceiling and in front of the animal, removing 43.8cm of the top of the enclosure and 72.4m of the front of the enclosure from the available space that the animals were able to move in (not depicted in figure 2).

A 5.08cm high by 4.45cm wide by 42.86cm long rectangular Styrofoam bar served as the obstacle. The bar was attached to a stepper motor which rotated the obstacle into and out of the path of the animal. The motor was attached to a belt linear actuator (Igus, RI) which moved the obstacle back and forth along the length of the treadmill. The whole obstacle apparatus was attached to the ceiling of the treadmill enclosure such that the top of the obstacle bar was 7.62cm above the treadmill floor. Additionally, a speaker was placed in the ceiling to play audio tones, and a small slot in the front of the enclosure allowed food rewards to be placed on the treadmill belt and carried to the animal. Treadmill speed, obstacle speed, and timing of the audio tones were measured using a hall-effect angular position sensor, stepper quadrature encoder, and electret microphone, respectively.

Eight cameras were positioned around the enclosure and captured video of the animals performing the tasks at 100Hz (SIMI Reality Motion Systems GmbH, Germany). Camera calibration was performed after each recording session to determine the positions and angles of each camera relative to each other, allowing for 3D triangulation of any markers that appear in at least two cameras. UV floodlights were placed around the recording room to enhance visibility of our UV reactive joint markers (see Kinematics). The cameras were synchronized to each other and to the neural data with TTL sync pulses. Additionally, 16 radiofrequency antennas were placed above and around the treadmill enclosure to receive the neural data transmitted through our wireless headstage (figure 2a).

The video capture was controlled through the Simi Motion software and the neural data capture was controlled through Blackrock Microsystem’s Central software. The treadmill, obstacle, and audio tone playback were controlled through a custom C++ program.

### 6.3 Behavioral Tasks

Animals performed either basic unobstructed locomotion by walking on the treadmill without any other interactions, or obstacle avoidance by stepping over an incoming obstacle during walking. Each of these tasks were carried out in blocks. Before entering the treadmill enclosure, animals were trained to enter a primate chair which allowed us to attach the wireless recording headstages (Cereplex W, Blackrock Microsystems, UT). The fur on the hind limb was shaved and the joint markers were also painted on at this time. Animals were then allowed to enter the treadmill enclosure where they were able to move freely.

During obstacle avoidance blocks, the treadmill was first turned on at 2.2 km/h. The obstacle would be in position in the front of the treadmill, but unmoving for the first minute to allow the animals to settle into a natural walking rhythm. At the start of each trial, a “go” tone was played to indicate that the obstacle was about to move. The operator would wait for a specific point in the gait cycle before starting the obstacle movement. The obstacle would move forward at 2.2 km/h until it was past the animal, and then rotate up out of the way of the animal. The obstacle was then moved back to the front of the treadmill and rotated down into the path of the animal to assume the starting position for the next trial. After stepping over the obstacle and while it was moving back into position, a “success” tone was played and a food reward was placed in the front of the treadmill which would be carried by the moving belt to the animal. After a few seconds to allow the animal to eat the reward and resume normal walking, the “go” tone would play again to initiate the next trial (figure 2b). At the end of the obstacle avoidance block, the treadmill would be turned off.

During basic walking blocks, the treadmill would be turned on and the animal would walk continuously without any obstacle or food interaction for 2-5 minutes. We would also include some of the strides at the beginning of the obstacle avoidance block before the obstacle was moved for the first time as basic walking trials. We excluded the first two strides after the treadmill was turned on and the last two strides before the treadmill was turned off to avoid any transition effects. All animals were trained to proficiently step over the obstacle without hitting it before experimental recordings were initiated. We recorded 38 obstacle avoidance trials from animal B and 43 trials from animal S, along with 49 basic walking trials from animal B and 66 trials from animal S.

### 6.4 Surgery

All surgical procedures were performed under general anesthesia induced through isoflurane. Animals were sedated with Ketamine (15 mg/kg) and Midazolam (0.05 mg/kg), and given Buprenorphone (0.01 mg/kg) pre-and intra-operatively. Buprenorphone SR (0.2 mg/kg) and Meloxicam (0.2 mg/kg) were adminsitered post-operatively. A craniotomy was performed and 96-channel multielectrode arrays (Blackrock Microsystems, UT) were inserted into leg area of primary-motor cortex (leg-M1) which was identified via cortical landmarks (figure 3a). Electrodes were platinum, 1.5mm in length, and attached to a percutaneous pedestal that was fixed to the skull. Animals were given at least one week to recover after the implantation surgery before resuming the behavioral tasks. Animal B experienced some motor deficits in the contralateral limbs initially after the surgery, although it was unclear whether the cause was from neurological damage during the implantation or poor positioning on the surgery table causing strain on the limb. However, he was able to fully recover walking ability within a week of walking on the treadmill post-surgery.

**Figure 3:**
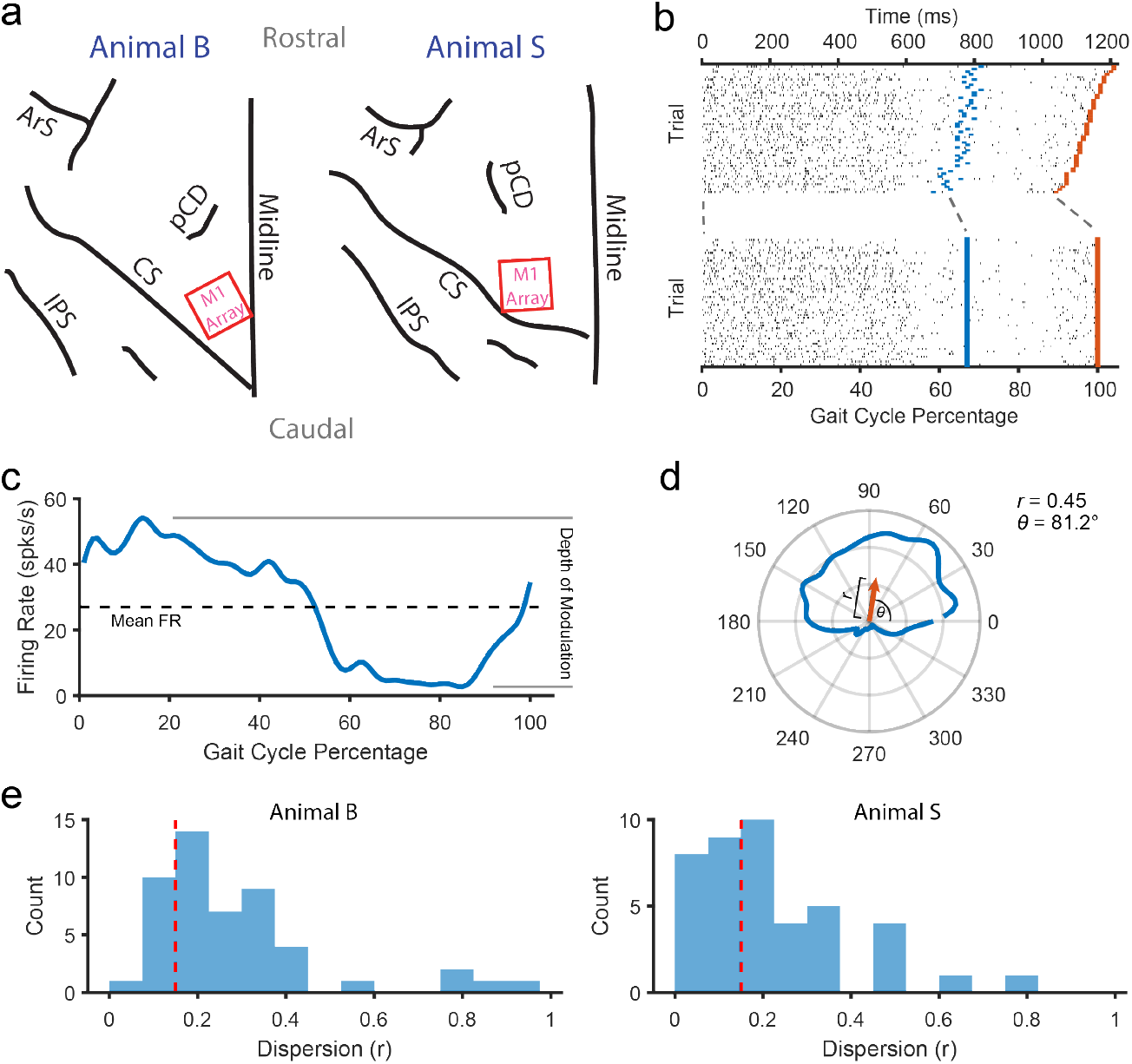
Neural recording and processing. **a** Microelectrode array implant locations. Red rectangles indicate where the Utah arrays were implanted in leg-area of M1 in the left hemisphere. *ArS* : Arcuate Sulcus, *pCD* : pre-Central Dimple, *CS* : Central Sulcus, *IPS* : Intraparietal Sulcus. **b)** Example raster of a neuron from animal B across multiple trials normalized to the gait cycle. Blue lines indicate foot-off time points (stance to swing transitions) and orange lines indicate foot-contact time points (swing to stance transitions). **c)** PETH of the neuron shown in **a** after normalization. Dotted black line indicates the average firing rate across the gait cycle, and depth of modulation is calculated as the amplitude of the firing rate across the gait cycle. **d)** The PETH in **c**, but shown as a polar plot. Circular statistics were used to calculate the average directional vector (orange arrow). The magnitude of the vector is the dispersion, *r*, of the neuron and the angle, *θ*, is the preferred phase of the neuron. **e)** Distribution of dispersion values for the whole recorded population of neurons for animal B (left) and animal S (right). Dotted red line indicates the cutoff value of 0.15 for classifying a “weakly modulated” neuron.

### 6.5 Kinematics

We used UV reactive colored body paint to mark the positions of 6 joints of the right hindlimb. We identified the joints through bony landmarks and painted circular markers over the iliac crest (crest), greater trochanter (hip), femur lateral epicondyle (knee), lateral malleolus (ankle), 5th metatarsal head (knuckle), and 5th distal phalanx (toe). Multi-camera video tracking was used to determine the 3D position of each of the joints (SIMI Reality Motion Systems, Germany). The direction of the treadmill movement was determined through markers placed on the side of the treadmill and the kinematics axes were rotated so that the x-axis corresponded to the direction of walking, the y-axis corresponded to the height, and the z-axis corresponded to medial-lateral movement. Because the animals were able to freely move back and forth along the length of the treadmill, we normalized the x-position of each of the joints to the x-position of the iliac crest.

Additionally, the timing of gait events (e.g. right hand off, left foot strike, ect.) were obtained manually by inspecting the video and marking the frame when the event occurred. A stride was defined as the time from one heel-strike of the limb to the next heel strike; the stance phase was defined as the period from the first heel-strike to the next toe-off, while the swing phase was defined as the toe-off to the next heel-strike.

### 6.6 Neural data processing

Intracortical recordings were obtained at 30kHz, band-pass filtered (300-3000Hz), and thresholded at 4x the standard deviation for spike events. Spikes greater than 1000 uV in amplitude were rejected as noise. For each channel, we used Wave Clus superparamagnetic clustering to semi-manually extract waveform templates (Quiroga et al., 2004), and used subtractive waveform decomposition for automated template matching of the thresholded spikes (C. Vargas-Irwin & Donoghue, 2007). We were able to isolate 50 neurons from animal B and 42 neurons from animal S. Spike counts were obtained by binning the number of spike events into 10ms bins corresponding to each frame of the video data. Single trial smoothed firing rate values were computed by convolving with a Gaussian kernel (s.d. 40ms).

Because each stride could vary in length, to compare across strides, we normalized the neural and kinematic data to the gait cycle. The start of the stride (heel-strike) was defined to be 0%, the toe-off frame of each stride was defined to be the average duty factor (67% for animal B and 69% for animal S), and the end of the stride (next heel-strike) was defined to be 100%. We used piecewise-linear time warping to interpolate the data at each percentage point from 0-100% (Matlab *interp1()* function). In general, the durations of the gait cycle were very consistent across trials (standard deviation was 44ms for animal B, compared to the stride duration of 1126ms and 65ms for animal S, compared to the stride duration of 1119ms), so we do not expect the normalization to have a significant effect on the analyses. Peri-event time histograms (PETHs) were computed by averaging the normalized spike counts across trials and then smoothed with a gaussian kernel (s.d. 20ms). Because of the unconstrained movement of the animals and wireless transmission of neural data, there would be periods where none of the antennas were able to receive the signal from the headstage. We excluded trials from analysis if more than 5% of the timepoints contained dropped signal.

### 6.7 Neural response characterization

Because the strides are cyclic, we utilized circular statistics to characterize the response profiles of each neuron (Berens, 2009; Drew & Doucet, 1991). The circular mean vector of the PETH was calculated by representing each value of the PETH as a polar vector and averaging across all percentge points (figure 3d). The preferred phase of the neuron was defined as the angle of the mean vector while the dispersion was defined as the magnitude. The depth of modulation was calculated as the difference between the maximum and minimum firing rate of the PETH, and the mean firing rate was calculated as the average firing rate across all the gait percentages. To identify neurons whose activity was uniform across the gait cycle, we applied the Rayleigh test at *α* = 0.05 with Bonferroni correction for multiple testing. We classified neurons as multi-modal if there were more than one peak that was greater than 50% of the depth of modulation for more than 10% of the gait cycle. Finally, we classified neurons as strongly modulated if they had a dispersion value greater than 0.15. All processing was done in Matlab with the Circ-stat toolbox.

To quantify the magnitude of change in the kinematics and neural activity during the obstacle stride, we calculated the Mahalanobis distance between the signal during the obstacle stride and the signal during the stride before any obstacle movement (three strides before the obstacle stride), which served as the baseline reference. At each gait percentage of the reference stride, we obtained the inter-trial distribution of the neural or kinematic activity in 15 dimensional space. For kinematics, this was the 3 spatial positions of the hip, knee, ankle, knuckle, and toe. For neural data, using the full 50 or 42 dimensional neural space resulted in singular matrix issues so we instead used the top 15 dimensions after performing PCA on the neural firing rates. After obtaining the mean and covariance matrix of this reference distribution, the Mahalanobis distance was computed for each trial of the obstacle stride at that gait percentage. The distance was calculated for all percentages of the gait cycle. The average and standard deviation of the distances across trials is what is shown in figure 5k-l. We also computed the cross-correlation between the average Mahalanobis distance of the neural activity and the average distance of the kinematics at various time lags ranging from -20 to 20 gait percentages. We used the Matlab functions *mahal()* and *crosscorr()* to implement the analyses.

**Figure 4:**
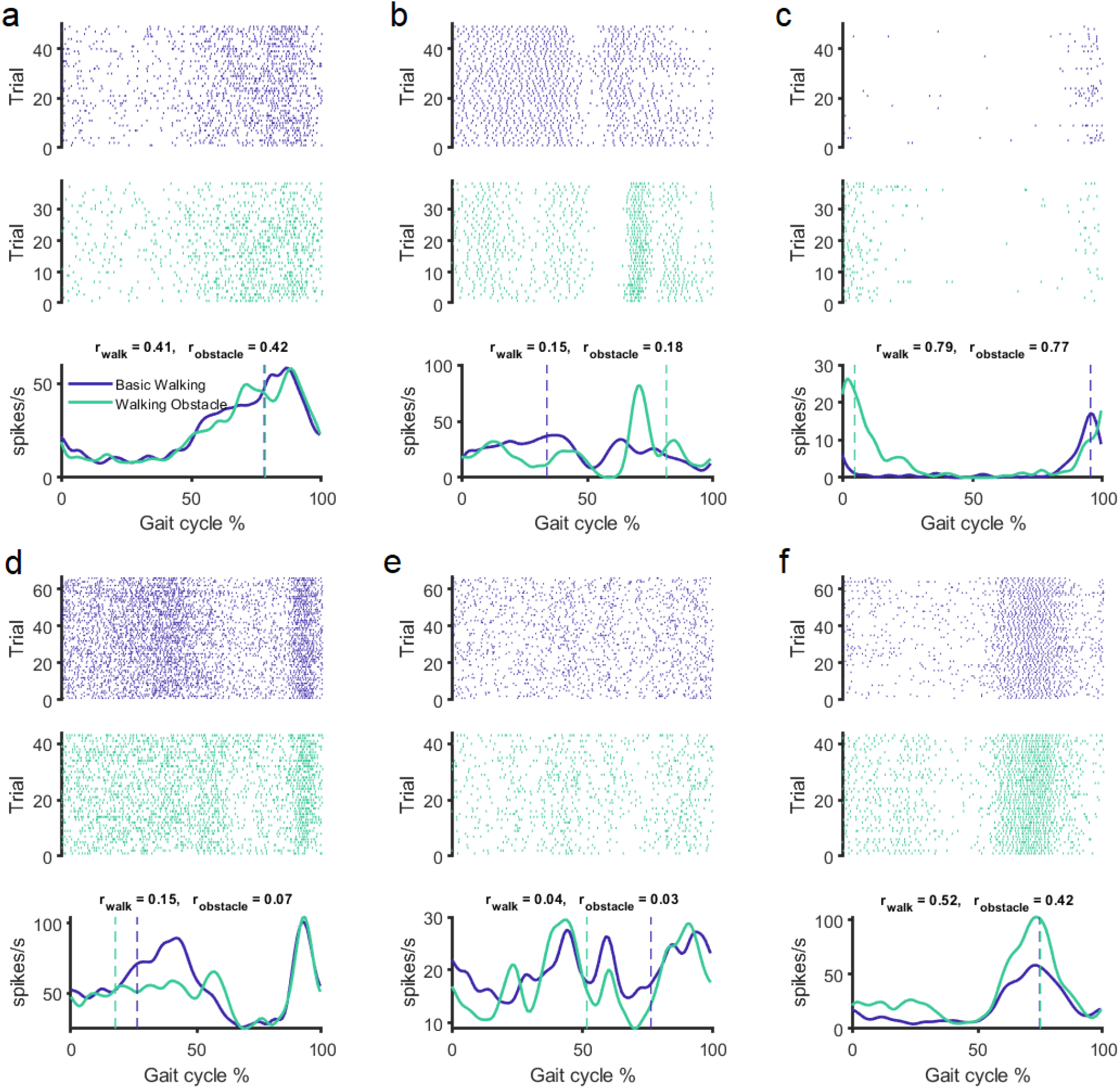
Example M1 neurons during basic walking and obstacle stepping. Example raster plots and PETHs from a M1-leg neurons. Top raster plot is for basic walking trials while bottom raster plot is for the obstacle stepping trials. r values represent the dispersion of the neural activity around the gait cycle, and the dotted lines represent the preferred phases of the neuron. **a-c)** Neurons from animal B, **d-f)** Neurons from animal S.

**Figure 5:**
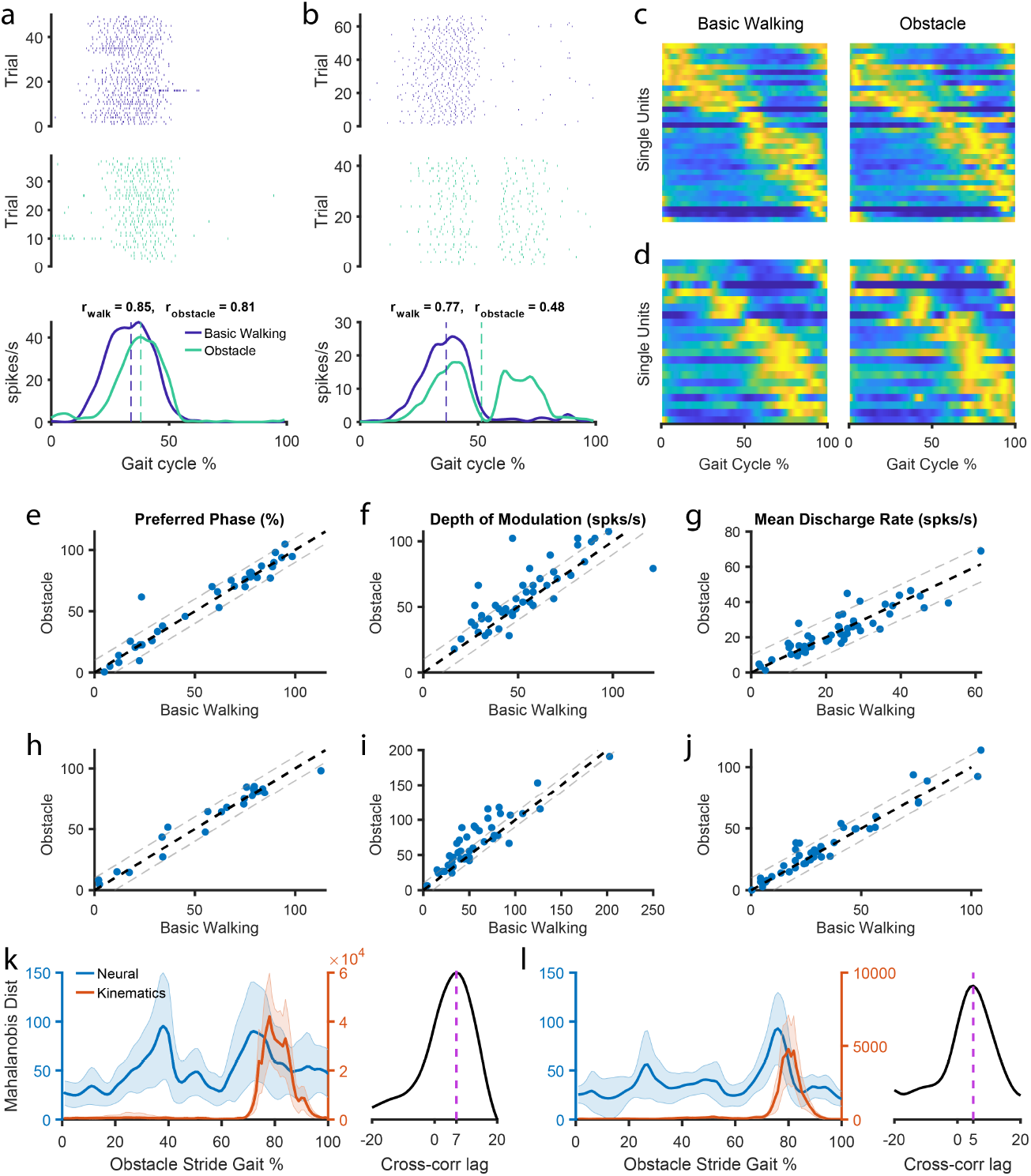
Most M1 neurons are phasically tuned to the gait cycle and increase their depth of modulation during obstacle stepping. **a)** Example plot of a neuron from animal B demonstrating a small shift in preferred phase. **b)** Same as in **a** but for an example neuron from animal S demonstrating complex changes in activity in response to obstacle stepping. **c)** Activity of all phasically modulated neurons for animal B during both basic walking (left) and the stride stepping over the obstacle (right). Neurons are sorted by ascending preferred phase of the basic walking trials, for both basic walking and obstacle stepping plots. The activity of each neuron is normalized to its maximum firing rate. **d)** Same as in **c**, but for animal S. **e**,**f**,**g)** Changes in preferred phase, depth of modulation, and average firing rate respectively of the neural population between the basic walking stride and the obstacle stride. Each point represents an individual neuron. Thick dotted line indicates no change, thin dotted lines delineate a change in 10% for preferred phase and 10 spks/s for depth of modulation and average firing rate. **h-j)** Same as in **e-g** but for animal S. **k)** Mahalanobis distance between the stride over the obstacle and the stride before any obstacle movement for the population of neural firing rates (blue) or the kinematic variables (orange). Right plot shows the cross-correlation between the neural and kinematic distances across multiple gait percentage lags with the dotted line indicating the peak lag. Positive indicates neural lagging kinematics. **l)** Same as in **k** but for animal S.

### 6.8 Demixed principle component analysis

We used dPCA to find task-specific subspaces within the population activity (Kobak et al., 2016). dPCA is similar to PCA in that they are both linear dimensionality reduction methods that projects high dimensional time series data into a lower dimensional space via a decoder matrix:

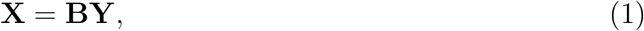

where **Y** is a *n* x *p* matrix representing the mean-subtracted PETHs of *n* neurons at *p* samples/datapoints;

**X** is the *m* x *p* matrix of the activity of *m* latent states, with m <n;

**B** is the *m* x *n* decoder matrix that projects each of the *n* dimensions to the lower *m*-dimensional space.

We refer to each of the *m* variables as a *latent dimension* or *neural mode*. However, unlike PCA which finds the projection that maximizes the variance accounted for by each of the *m* dimensions regardless of any task-related parameters, dPCA attempts to find subspaces that are related to these parameters. Kobak et al. provides an in-depth walkthrough of the dPCA algorithm and implementation. We applied the Matlab toolbox supplied by Kobak et al. for all our dPCA calculations.

For our data, we chose to use two task parameters: gait cycle percentage, (which in the unnormalized case, would correspond to time), and stride type. Gait cycle percentage varied from 0-100% and stride type varied from three strides before the obstacle stride to two strides after. We combined stride type and stride-percentage interaction terms into the stride type marginalization, since we expect the stride-related neural activity to also be time-varying. Therefore, the gait percentage marginalization subspace, which we call the stride-invariant subspace, should only vary with the gait cycle percentage and not with the stride type, while the stride marginalization subspace, which varies with both gait percentage and stride type, will be our stride-dependent subspace. dPCA requires a dimensionality of the model be specified explicitly, so we chose a dimensionality of 10 (5 for the stride-invariant and 5 for the stride-dependent subspace) based on findings of previous studies, but we also tested our results for dimensionality of 6 and 14 (split equally between the two subspaces), and did not find any significant changes on our results.

Additionally, we wanted to test which aspects of the stride-dependent signals truly correspond to changes in neural activity due to the obstacle, and which aspects are just random fluctuations due to noise. To get an estimate of the noise distribution, we performed dPCA using just the first stride and the last stride. These strides occur before the obstacle begins to move, and two strides after the obstacle has been traversed, respectively, and are essentially the same as basic walking strides. Therefore, we do not expect any activity related to a volitional gait adjustment movement to be present in the neural activity, and thus any signals found in the stride-dependent subspace using only these strides would be just noise. We took these values as the null distribution, and used a conservative cutoff value of the 99.99th percentile of this noise distribution to determine which components of the signals in the full dPCA analysis are significantly greater than noise.

Finally to quantify the timing of the changes in the stride-dependent subspaces in relation to the changes in the movement, we used the same analysis as before during the neural response characterization (figure 5k-l). We calculated the cross-correlation between the mean Mahalanobis distance of the kinematics in the obstacle stride and the first component of the stride-dependent subspace in the obstacle stride at varying time lags.

### 6.9 Principle Angles

Within each marginalization subspace, the dimensions are orthogonal (that is, the rows within each encoder matrix form an orthonormal basis). However, because each marginalization projection is calculated independently, the resulting subspaces do not have to be orthogonal to each other, meaning there could be a significant amount of overlap between the subspaces. To quantify the alignment between two subspaces, we calculated the principle angles between our stride-invarient (**E**_*inv*_) and stride-dependent (**E**_*dep*_) subspaces (Gallego et al., 2018). The angles range from 0° to 90° which indicates perfectly overlapping or perfectly orthogonal subspaces, respectively. To obtain the principle angles, we performed singular value decomposition on the product of the two encoder matrices:

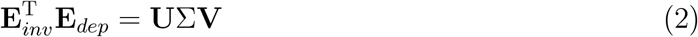

The diagonals of the Σ matrix are the cosines of the principle angles, ordered from smallest to largest.

We performed dPCA on both the neural PETHs as well as the trial-averaged kinematics for the obstacle avoidance trials, and calculated the principle angles between the **E**_*inv*_ and **E**_*dep*_ for both. To get a sense of the sampling variance, we performed 500 bootstraps on the trials and calculated the prinpcle angles for each bootstrap resample. The 95% confidence intervals of the bootstrap distribution are shown in figure 6d,i.

**Figure 6:**
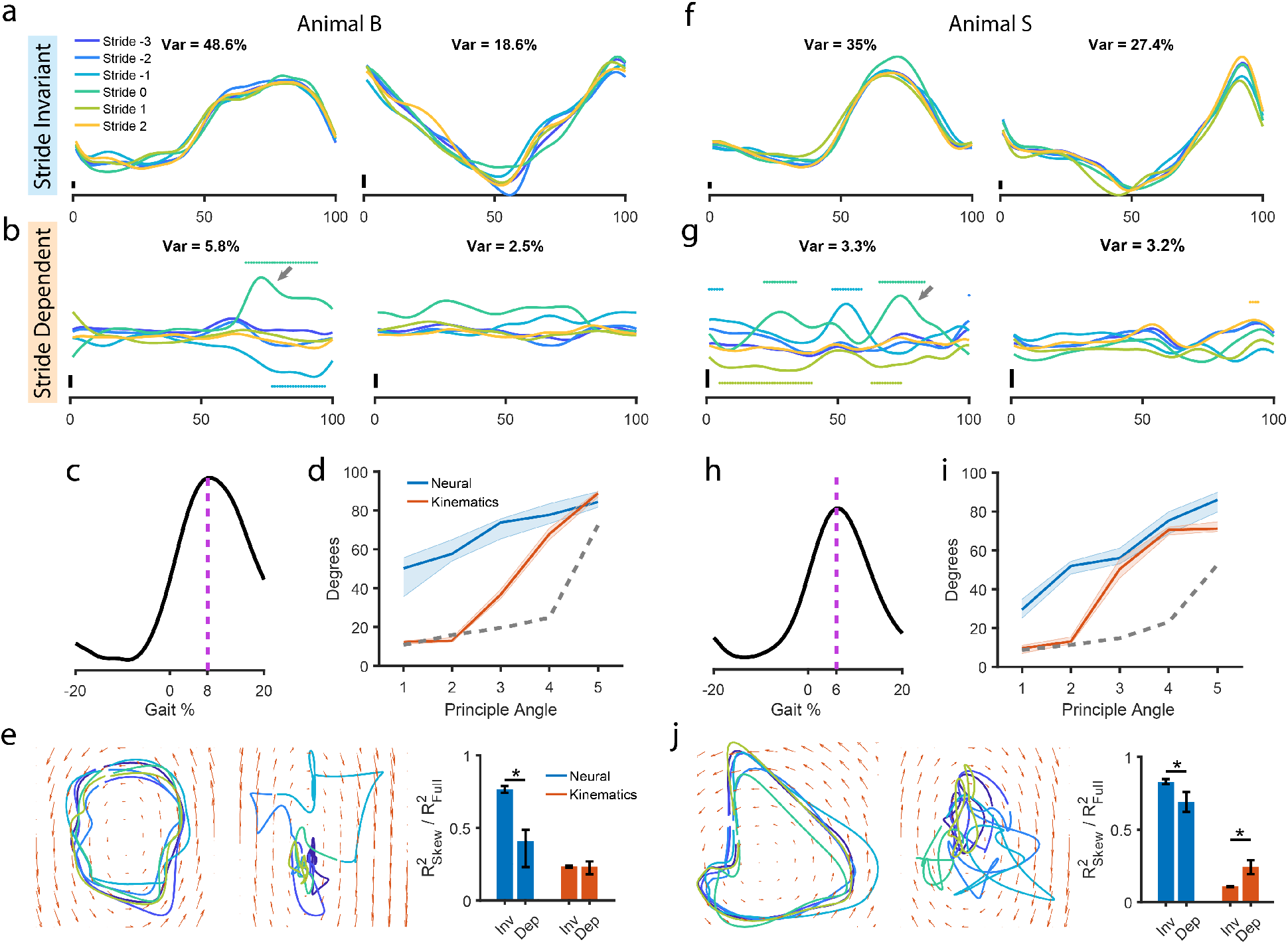
dPCA separates neural activity into obstacle stride-invariant and stride-dependent subspaces. **a-b**,**f-g)** Top two stride-invariant (**a**,**f**) and stride-dependent (**b**,**g**) demixed components for animal B. Each of the strides surrounding the obstacle stride (Stride 0) are plotted individually. Numbers at the top represent percentages of the total variance for each component. Arrows indicate the increase in the stride-dependent neural modes during the obstacle stride that account for the peaks in **c** and **h**. Colored dots indicate points where the stride-dependent components were significantly greater than noise (see Methods).**c**,**h)** cross-correlation between the change in kinematics and the change in activity within the stride-dependent subspace across multiple gait lags, similar to figure 5**k,l. d,i)** Principle angles between the stride-invariant subspace and stride-dependent subspace for both neural dPCA and kinematics dPCA. Error bars represent 95% confidence intervals for 500 bootstrap resamples. Dotted grey line represents the 97.5th percentile of the null distribution of completely overlapping subspaces. **e**,**j)** First two stride-invarient components (left plot) or stride-dependent components (center plot) are plotted against each other to visualize rotational structure (or lack thereof). The slope fields for the rotations inferred with jPCA are shown as orange arrows. Right plot: ratio of the jPCA model R^2^ to an unconstrained LDS model R^2^ to quantify the strength of rotations. For dPCA used on neural activity, maximum tensor entory (MTE) was used to generate surrogate datasets and the R^2^ ratios was also calculated for the stride invariant subspace of the surrogates. Error bars are 95% confidence intervals for 500 bootstrap reshuffles. Stars indicate statistically significant difference (Wilcoxon rank sum test, *α* = 0.05).

We hypothesized that the stride-invarient and stirde-dependent subspaces are separate. To build a distribution of the principle angles for the null-hypothesis, we used two subspaces that we know are completely aligned other than from inter-trial variance: the **E**_*inv*_ ‘s (or **E**_*dep*_’s) from two different bootstrap resamples. We performed two bootstrap resamples and calculated the principle angles between **E**_*inv, resample1*_ and **E**_*inv, resample2*_ as well as the angles between **E**_*dep, resample1*_ and **E**_*dep, resample2*_. This was repeated 250 times to obtain our null distribution. The 97.5th percentile of the principle angles for this null distribution is shown in figure 6d,i.

### 6.10 Rotational structure

To model rotational dynamics within the subspaces, we employed jPCA which is a specialized variant of a linear dynamical system (LDS) (Churchland et al., 2012). The underlying assumption of dynamical systems models is that the current state of the latent components is predictive of future states. LDS models the relationship between states as a difference equation with time-evolution matrix **A**:

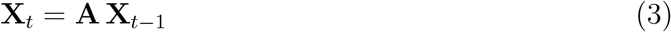

Here, **X**_t_ is the low-dimensional neural activity in the stride-invariant or stride-dependent subspace at gait percentage *t*. In general-form LDS, **A** is unconstrained and can be obtained analytically through least-squares regression. In jPCA, **A** is constrained to be skew-symmetric which results in a dynamical system that contains solely rotations. While it is possible to solve for **A** analytically in the constrained case, we followed the algorithm implemented in Churchland et al. which uses a gradient-based optimization method. After obtaining **A**, one can visualize the rotational dynamics by displaying the slope fields of the difference equation in the top two dimensions, as in figure 6e,j. The jPCA algorithm was implemented with the Matlab code provided by Churchland et al. Note that because we are already in a low-dimensional space, we did not perform the pre-processing step of using PCA to project into 6 dimensions before fitting for **A**, as was done in Churchland et al.

To quantify the strength of rotational structure in each of the subspaces, we calculated the ratio of the jPCA fit R^2^ to an unconstrained LDS fit R^2^. The unconstrained LDS can contain both rotational and non-rotational dynamics, but if the dynamics in the data are purely rotational, then the best unconstrained LDS **A** would be the same as the jPCA **A**, and the ratio would be 1. The more non-rotational dynamics are present, the more divergent the jPCA fit will be from the best LDS fit and the lower the ratio. We calculated this metric for the stride-invariant data and the stride-dependent data for both neural and kinematic dPCA.

### 6.11 Poisson linear dynamical system model

We employed an unsupervised dimensionality reduction approach to complement our dPCA analysis. Like PCA, PLDS assumes that the recorded neural activity is the result of underlying latent states (Aghagolzadeh & Truccolo, 2016; Macke et al., 2011; Xing et al., 2019). However, in PLDS the spiking of each neuron is modeled by a (conditionally) Poisson process rather than the usually assumed Gaussian distribution. Additionally, PLDS explicitly models temporal dynamics by incorporating a (Gaussian) linear dynamical system similar to what was discussed in the previous section. The full model is described by:

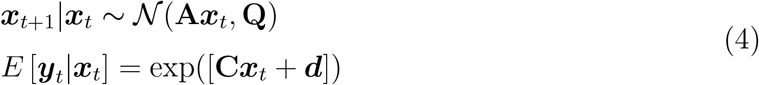

where ***y***_*t*_ is the vector of recorded spike counts of all neurons at time point *t*,

***x***_*t*_ and ***x***_*t+1*_ is the vector of the activity of the latent neural modes at time *t* and *t+1* respectively,

**C** is the matrix of weights relating the neural modes to the conditional intensity function of the neurons, analogous to the **B** in dPCA,

***d*** is the baseline firing rate of the neurons,

**A** is the time evolution matrix governing the temporal dynamics of the neural modes, analogous to the **A** matrix of the unconstrained LDS model in the previous section,

**Q** is the covariance of neural modes after the time evolution, i.e. the covariance of additive Gaussian noise.

Unlike PCA and dPCA, there is no analytic solution for the model, so we used the expectation maximization algorithm to infer the latent state activity and model parameters. Also, due to the non-linearity, there is no closed-form solution to finding the posterior probability of the latent neural modes given the spiking activity observations and estimated parameters in the E-step, so we employ the Laplace approximation which formulates the latent state posterior density at each time step as a Gaussian conditioned on the corresponding observations. We then estimate the values of the neural modes as the mean of this Gaussian which is calculated for each time point of each trial. A detailed description of the inference algorithm can be found in (Aghagolzadeh & Truccolo, 2014; Macke et al., 2015). We ran the EM algorithm for a total of 60 iterations.

To estimate the dimensionality of the underlying neural subspace, we employed the Akaike information criterion (AIC):

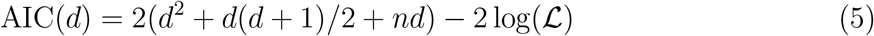

where *d* is the number of neural modes and *n* is the number of neurons and *L* is the likelihood function. The value that minimized the AIC was chosen as the final model dimensionality. Using this metric, we estimated the best-fit model dimensionality to be approximately nine.

We additionally created a model dataset of neural activity that was constructed from the kinematics, following the method of (Gallego et al., 2018). We simulated as many neurons as there were in our recorded dataset for each subject. Each neuron was modeled as a Poisson process with a *λ* defined as the weighted sum of the kinematics:

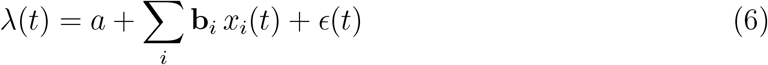

where x_i_(t) is the value of the normalized ith kinematic variable, a is a baseline firing rate randomly sampled from [0 0.1], b_i_ is the weight of the ith kinematic variable onto the neuron, drawn from a Gaussian distribution of mean 0 and variance 1, and *ϵ* is additive Gaussian noise with zero mean and a variance of 0.01. The kinematics consists of the 3D coordinates of the 5 joints, and were normalized by subtracting the mean and scaling the amplitude to 1. The *λ*s were scaled so that they were all positive. We then generated spike trains for each of the neurons and ran the PLDS analysis on the simulated neural data.

### 6.12 Spread metric

The PLDS trajectories shown in figure 7 were created by time-normalizing the activity in the latent states of each trial to 0-100% of the gait cycle and then averaging across trials. We displayed the first three PLDS dimensions to visualize the time-varying activity of the neural modes throughout the 6 gait cycles as the animal stepped over the obstacle. Within this three-dimensional space, certain 2-dimensional projections resulted in large amounts of overlap in the neural trajectories across the different strides. We used the Matlab function *viewmtx()* to calculate the projections at various azimuth and elevation viewing angles. We then chose the projection that contained the most overlap as our stride-invariant subspace. A different viewing angle which highlights the changes in the neural trajectories was subjectively chosen as the stride-dependent subspace.

**Figure 7:**
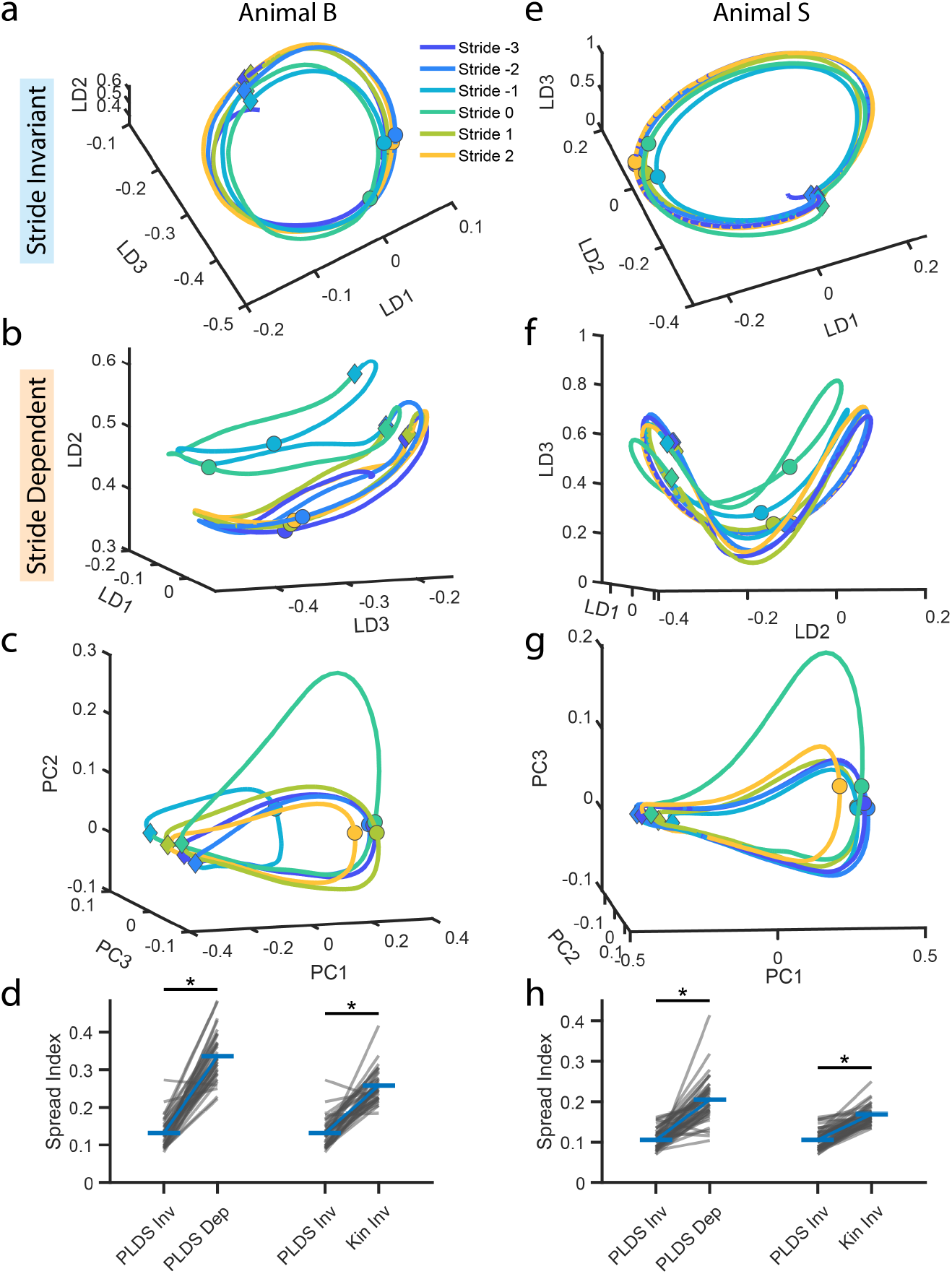
PLDS also extracts neural modes invariant to obstacle stepping. **a**,**e)** Neural trajectories from three neural modes, or latent dimensions (LD), inferred from the PLDS model. Each of the strides surrounding the stride over the obstacle (Stride 0) are plotted individually. The view angle of the trajectories is chosen to maximize the overlap between all of the strides according to a spread index (see methods). The projection into this view represents the stride invariant subspace. Circular dots indicate the transition from the stance phase to the swing phase in each stride, while diamond dots indicate the transition from the swing phase to the stance phase of the next stride. **b**,**f)** Same plot as in **a** and **e** but with a view angle which highlights the differences across strides, representing a stride dependent subspace. **c**,**g)** Top three PCA components of the kinematics for the same strides as the PLDS plots. **d**,**h)** Spread index quantifying how much the trajectories overlap or diverge across strides (see methods). We compared the amount of divergence between the trajectories in the stride invariant neural subspaces with the stride dependent neural subspaces (left) as well as the stride invariant neural subspaces with the stride invariant kinematic subspaces (right). Stars indicate statistical significance (Wilcoxon signed rank test, *α* = 0.05). **a-d**: animal B, **e-h**: animal S

To quantify the amount of overlap in the low-dimensional PLDS projections between different strides, we created a spread metric which determines how divergent the neural trajectories become relative to a reference baseline stride. Because we expect the obstacle stride to have the greatest divergence from the other strides, we chose that stride as the baseline.

Let the vector ***a*** _ref,t_, be the coordinates of the reference stride in the projected 2D space at gait percentage *t*. For each of the other strides *i* = 1…5, we calculated the euclidean distance of the closest point to ***a*** _ref,t_ within 10 gait percentages,

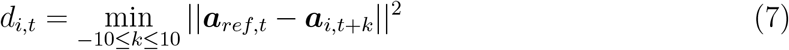

We chose to use all points within 10 gait percentages to account for any “slippage” due to imperfect time-normalization between the strides. We then took the maximum distance across all of the strides, max *d*_*i,t*_ to find the spread metric at gait percentage *t*. Finally, we obtained the final spread metric by taking the 90th percentile across gait percentages 20 *≤ t ≤* 80. We excluded the first and last twenty gait percentages to avoid artifacts from edge effects and used the 90th percentile in order to exclude outliers.

### 6.13 Variance calculations

For PCA and dPCA, we measured how much neural variance the low-dimensional neural modes were able to explain by calculating the variance acounted for (VAF):

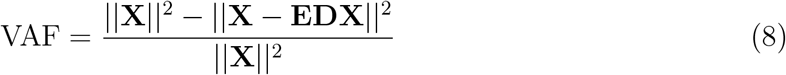

Where **X** is the matrix of PETHs of all neurons, **E** and **D** are the encoder and decoder matrices respectively, and the indicated norm is the Frobenius norm.

### 6.14 Decoding

We utilized a Wiener decoder of order 10 to predict kinematics at each gait percentage point, denoted by ***y*** _t_, using various neural inputs.

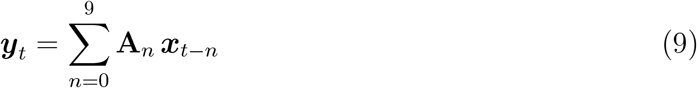

Here ***x*** _t-n_ is the vector of input features time-shifted by *n* gait percentages and **A**_n_ is the matrix of decoding weights, which we computed through least-squares regression on training data. For all of our decoding analyses, we used the toe height as the decoded kinematic variable.

We tested whether a decoder trained to predict kinematics during basic locomotion could generalize to obstacle avoidance movements. We used the firing rates of the full neural population as the input vector, and calibrated the decoder weights using the neural data and kinematics from the stride three strides before the obstacle stride (stride -3). We then used those weights to predict the kinematics during the subsequent strides, including the obstacle stride. We also tested the opposite, whether a decoder trained on the obstacle avoidance stride could decode kinematics during basic locomotion. In this case, the weights were trained on the neural and kinematic data during the obstacle stride and applied to each of the other strides. Note that for testing decoder generalization, we did not need to employ cross-validation since the training set for the decoder was completely separate from the testing set. However, we also measured how well the decoders trained on either the basic walking or the obstacle stride could decode the kinematics during that same stride. In these cases, we employed leave-one-out cross validation by removing one trial from the training set and using it as the testing set. As a control, we built a decoder trained on all of the strides, and calculated the decoding performance across all of the strides using leave-one-out cross validation as well.

When testing decoder performance using low-dimensional neural modes as inputs, we utilized the same procedure, but instead of using the neural firing rates as ***x*** _t_, we used the neural trajectories obtained with dPCA. We measured the decoding performance using just the latent dimensions within the stride-invariant subspace, or the dimensions in both the stride-invariant and stride-dependent subspace. In both cases, all of the strides were included in the training set, and leave-one-out cross validation was employed.

We measured decoder performance by calculating the mean squared error between the decoded kinematics, 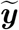, and the real kinematics across all gait percentages:

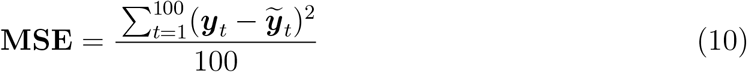

## 7 Results

### 7.1 Animals carried out volitional gait adjustments in an obstacle avoidance paradigm

Animals were successfully trained to enter the plexiglass enclosure on top of the motorized treadmill and continuously walked at 2.2 km/h. Within the enclosure, an actuated Styrofoam bar moved towards the animal at 2.2 km/h, giving the perception of an obstacle approaching along the treadmill belt (figure 2a). Both animals successfully raise their limbs to step over the obstacle without hitting it while maintaining their ongoing walking movements. A “go” tone would alert the animal of the oncoming obstacle, and the operator would manually time the start of the obstacle movement to a particular gait event (e.g. the instant the right hand made contact with the floor) in order to maintain consistency in the avoidance movement across trials. Although there was some amount of jitter in the timing of the obstacle initiation relative to the gait cycle, the variance was rather small (standard deviation: 2.7% of the gait cycle for animal B, and 2.8% for animal S). Each trial consisted of one obstacle run which included the stride where the leg was lifted over the obstacle (denoted as stride 0), three strides before the obstacle stride (strides -3 to -1), and two strides after the obstacle stride (strides 1 to 2, figure 2c). Additionally, we collected data from the animals as they continuously walked on the treadmill without any obstacle movement, which we will refer to as basic walking. Example neural activity and kinematics of an obstacle stepping trial, along with the gait pattern and obstacle position are shown in figure 2c.

### 7.2 Stepping movement form is preserved during obstacle avoidance while the step height is increased

The limb movement throughout all the strides, including the obstacle stride, follows the same stepping pattern. Horizontal and vertical positions of the joints increased and decreased at similar phases of the gait cycle, while the main difference between the obstacle stride and the other strides is in the magnitude of the movements. Both animals had to raise their hindlimbs considerably higher than during their normal walking gait in order to clear the obstacle (figure 2d-e,g-h, movie 1, p <1e-5, wilcoxon rank sum test). During basic walking, the average stride height was 3.66cm for animal B and 4.45cm for animal S, and increased to 17.95cm and 11.28cm for the obstacle stride, respectively. However, the stride durations were similar for all the strides, with the exception of the stride immediately before the obstacle stride in animal B, which was shorter than usual (figure 2f,i, p <1e-5, wilxocon rank sum test). In both animals, the right limb was the trailing limb over the obstacle and they had similar stride duty factors, 67% for animal B, and 69% for animal S.

The obstacle did not start moving until two strides before the obstacle stride in animal B and one stride before the obstacle stride in animal S, meaning that from the perspective of the animal, the first gait cycle in the obstacle avoidance trials should be essentially the same as the gait cycle during basic walking. Indeed, the stride height and duration are virtually identical between these two strides (figure 2e-f,g-h). Between the next two strides, the animals could see and were aware of the obstacle moving towards them, although the stride height remained unchanged for these strides. After clearing the obstacle, the stride height returns to normal pre-obstacle ranges. In summary, despite the large change in kinematics in the obstacle stride, the subjects returned to normal walking quickly after avoiding the obstacle, and with the exception of animal B taking a smaller and quicker stride right before the obstacle stride, they did not drastically alter their gait leading up to the obstacle stride.

### 7.3 M1 neurons show increased activity but little phasic reorganization during obstacle avoidance

We then characterized the response properties of individual neurons in leg area of M1. Figure 5a, b illustrates the activity of two example neurons during both basic walking (purple) and the stride over the obstacle (teal). In order to compare the activity across trials, which had some slight variations in duration, the spike trains were time-normalized to 0-100% of the gait cycle (methods, figure 3. Consistent with previous M1 recordings during locomotion, the majority of neurons tended to increase their firing rate during specific phases of the gait cycle (Foster et al., 2014; Xing et al., 2019; Yin et al., 2014). Although many of these neurons fired at around a single phase of the gait cycle (e.g. figure 5a, figure 4c), we also found neurons that had multimodal peri-event time histograms (PETHs) (e.g. figure 4d). We utilized circular statistics (Berens, 2009; Drew & Doucet, 1991) on the unimodal PETHs to determine the preferred phase and dispersion (*r*) of the neurons around the gait cycle (figure 3d). We applied the Rayleigh test to identify neurons which were not significantly modulated to the gait cycle and further classified unimodal, modulated neurons as weakly modulated if they had a r value less than 0.15 during basic walking. Figure 3e shows the distribution of dispersion values and indicates that most of our recorded neurons are strongly modulated. Since preferred phase is only well defined for neurons that are both significantly modulated to the gait cycle and have a unimodal PETH, we excluded neurons that are multimodal or weakly modulated from preferred phase calculations (but not depth of modulation or mean firing rate calculations).

We found that for both animals, the preferred phases of the population of recorded neurons spanned the whole gait cycle (figure 5c,d), consistent with previous studies in felines and macaques (Beloozerova & Sirota, 1998; Drew et al., 2008; Foster et al., 2014; Xing et al., 2019; Yin et al., 2014). Additionally, neurons tend to fire at the same phase during the obstacle avoidance step as during basic walking (figure 5c-e,h). Only 4/34 (animal B) and 2/22 (animal S) of the strongly modulated neurons had shifts in preferred phase greater than 10% of the gait cycle. However, most neurons tended to increase their depth of modulation during the obstacle step (average increase of 6.37 spks/s for animal B, 11.62 spks/s for animal S), reflecting the larger amplitude of the movement (figure 5f,i). The mean firing rates also saw an overall increase, although the change is not as large (average increase of 0.52 spks/s for animal B, 2.25 spks/s for animal S, figure 5g,j). Finally, we note that many of the neurons exhibited complex changes in firing activity beyond simple phase shifts or changes depth of modulation (e.g. figure 4b,d).

Lifting the leg over the obstacle involves a decision to move the limbs in a specific way, which requires descending input from motor cortex. We therefore hypothesized that there would be a change in neural activity preceding the modifying movement to the gait. We determined when the changes in the obstacle avoidance stride occurred by calculating the Mahalanobis distance between stride 0 (the obstacle stride), and stride -3 (the stride before any movement in the obstacle) at each percentage of the gait phase. Unsurprisingly, we saw a large increase in the difference between the kinematics during the the swing phase (figure 5k,l orange plot, p <1e-5, wilcoxon rank sum test), corresponding to the increase in step height. We also observed an increase in the difference of the neural population immediately before the kinematic divergence (figure 5k,l blue plot, p <1e-5, wilcoxon rank sum test), giving evidence for the presence of an efferent signal in M1 related to the action of lifting the leg over the obstacle. Cross-correlation analysis determined that the neural modulation preceded the change in kinematics by 7% of the gait cycle in animal B and 5% in animal S, which, using the average stride durations, corresponds to 72.1ms and 52.3ms respectively. One possible explanation for this signal could be that cortex engages separate sub-populations for maintaining locomotion and for carrying out voluntary gait adjustment, and would only recruit the neurons related to voluntary movement during the obstacle stride. This may be the case for the whole cortical population of neurons, however, within the population we are recording from, we did not see the emergence of any clusters in the upper left in our depth of modulation plots, indicating the lack of obstacle-avoidance specific neurons (figure 5f,i).

Taken together, these results indicate that M1 neurons are active during basic locomotion and that the same population is also predictive of volitional gait-adjusting movements. Previous work in rodents have shown that M1 population activity, although present during both basic treadmill walking and stationary voluntary movements, resides in separate subspaces across the two behaviors (Miri et al., 2017). We wanted to determine whether this compartmentalization of neural activity is maintained when both types of movements have to be carried out simultaneously such as in obstacle avoidance. We therefore employed dimensionality reduction techniques in order to investigate whether M1 utilized specific neural subspaces for carrying out the gait adjustment, and whether these subspaces are distinct from the ones present during basic locomotion.

### 7.4 dPCA reveals division of neural modes into obstacle-related and obstacle-invariant subspaces

Demixed principle component analysis (dPCA) is a statistical method which decomposes neural activity into subspaces associated with specific task parameters. For example, in a vibrotactile discrimination task, it was able to isolate neural activity related to the type of presented stimulus, the decision of the subject, and also activity invariant to either of those parameters (Rossi-Pool et al., 2017). Here, we used dPCA to extract subspaces that were agnostic to the gait adjustment (stride-invariant) and also subspaces which captured the change in population activity during the gait adjustment (stride-dependent). We found that 10 dPCA components were able to explain 87.2% and 88.4% of the variance for animal B and animal S, respectively.

The stride-invariant subspace accounted for most of the neural variance (77.1% for animal B, 80.7% for animal S), and indeed showed very little modulation across strides (figure 6a,e), preserving the same time-varying activity throughout all the strides during obstacle stepping. The stride-dependent activity accounted for 11.9% of the variance in animal B and 11.0% for animal S. We observed large deviations in the stride-dependent components during the stride immediately before and the stride over the obstacle (figure 6b,f, p <0.0001, see methods), suggesting that these components capture an increase in M1 engagement during those strides. Additionally, we compared the timing of these shifts in relation to the changes in the kinematics during the swing phase and found that, like with the population firing rates, the activity of the stride-dependent components preceded the gait changes, (figure 6c,h).

Although dPCA was able to find subspaces that are invariant and subspaces that are dependent on the gait modification, the nature of the algorithm does not guarantee that the stride-invariant and stride-dependent subspaces are orthogonal. For example, we were also able to find stride-invariant and stride-dependent subspaces for the kinematics, yet these subspaces could be largely overlapping. Principle angles have been used in previous studies to determine the alignment between two subspaces Gallego et al., 2018. We calculated the principle angles between the stride-invariant and stride-dependent neural subspaces, and found that they were greater than the principle angles between the kinematics subspaces (figure 6d,i; methods) as well as a boot-strapped null distribution (methods, *p* <0.05 significance level).

It is also possible that these dPCA components reflect the activity of specific sub-populations of neurons, rather than the whole population. We investigated this possibility by examining the projection weights of each neuron (Gallego et al., 2018; Kobak et al., 2016). We found that the distribution of the weights for all of the components is unimodal, with small weights for most neurons and no outliers. Additionally, we performed cluster analysis on the neural weights (Kobak et al., 2016; Rodriguez & Laio, 2014) and found that the neurons formed only a single cluster, indicating that the dPCA components are distributed across the whole neural population rather than corresponding to specific sub-populations.

Finally, we hypothesized that since the cyclic locomotor rhythm was a common factor across all of the strides, the stride-invariant subspace would contain significant rotational dynamics, while the dynamics of the stride-dependent subspace, which represents the precisely timed motor adjustments on the gait, would be less rotational. We used jPCA to fit a rotational dynamical system to both sets of subspaces, and quantified the rotational strength as the ratio of the coefficient of determination (R^2^) of the jPCA model to the R^2^ of an unconstrained dynamical systems model (Churchland et al., 2012). Figure 6e,j illustrates the fitted rotational dynamics of the first two latent dimensions in both subspaces. In agreement with our hypothesis, we found that there was significantly more rotational structure in the stride-invariant than the stride-dependent subspace (two-tailed Wilcoxon rank sum test, *p* <1^-100^ for both animals, n = 500 bootstrap resamples for all dPCA statistical tests). In contrast, when we tested the kinematics, we found that the stride-dependent subspace had greater rotational structure (*p* <1^-100^ for animal S), or no difference in the rotational strength between the stride-invariant and stride-dependent subspaces (*p* = 0.9099 for animal B). Finally, although we chose to use a dimensionality of 10 in our dPCA model (5 dimension for stride-invariant, 5 for stride-dependent subspaces) based on previous dimensionality reduction studies (Sadtler et al., 2014; C. E. Vargas-Irwin et al., 2015; Xing et al., 2019; Yu et al., 2009), we tested these findings across a range of dimensionalities and obtained similar results.

### 7.5 Unsupervised dimensionality reduction shows similar separation of neural subspaces

dPCA is a powerful technique which uses labeled data to extract task-related subspaces. To ensure that our findings are not simply the result of the partially supervised nature of dPCA, we also employed an unsupervised dimensionality reduction model completely agnostic to the stride type. One such model, Poisson linear dynamical systems (PLDS), explicitly infers the low-dimensional dynamics through a time-evolution matrix, resulting in smooth single trial neural trajectories without the need for trial averaging (figure 7a-b,e-f)(Aghagolzadeh & Truccolo, 2014; Macke et al., 2011; Xing et al., 2019). We have previously shown that the PLDS neural modes were able to reconstruct limb kinematics just as well as the full population rates and better than PCA components during locomotion (Xing et al., 2019).

The top three latent components accounts for most of the variance (60.8% for animal B, 86.4% for animal S).The neural trajectories in these components for each of the strides are shown in in figure 7a-b for animal B and e-f for animal S. Interestingly, there are projections of this space where the neural trajectories completely overlap, regardless of the stride (figure 7a,e), while other projections reveal a divergence of the trajectories during the obstacle stride (figure 7b,f). We defined the stride-invariant subspace as the projection which results in the greatest amount of overlap across all of the strides. We quantified the amount of overlap using a spread metric (methods), and optimized for the projection which minimized this spread metric. There are two notable features in these projections that are consistent in both animals. First is the clear rotational structure in this subspace, corroborating our results from the dPCA analysis. Second is the high amount of invariance between the trajectories across the different strides, despite the large change in movement during the obstacle stride. We emphasize that unlike dPCA, the PLDS model is not designed to specifically identify neural modes invariant to any particular variable, so it is entirely possible that the modulation in response the the voluntary gait adjustment pervades all the top PLDS neural modes (e.g. as in figure 1c). Indeed, the top three principle components of the kinematics demonstrate this case, as there are no projections resulting in the same amount of overlap between the obstacle stride and the basic walking strides (figure 7c,g). When comparing smallest achievable spread index for the kinematics and the PLDS trajectories, the kinematics was significantly larger (one-tailed Wilcoxon signed rank test, n = 38 trials, *p* = 4.03×10^−8^ for animal B, n = 43 trials, *p* = 6.23×10^−9^ for animal S) by about 2.13 times for animal B and 1.58 times for animal S (figure 7d,h). As an additional control, we simulated a neural dataset using only kinematics data, following the method of (Gallego et al., 2018). Like with the kinematics PCA, there is no projection in which the ongoing activity is unaffected by the obstacle movement (figure 8).

**Figure 8:**
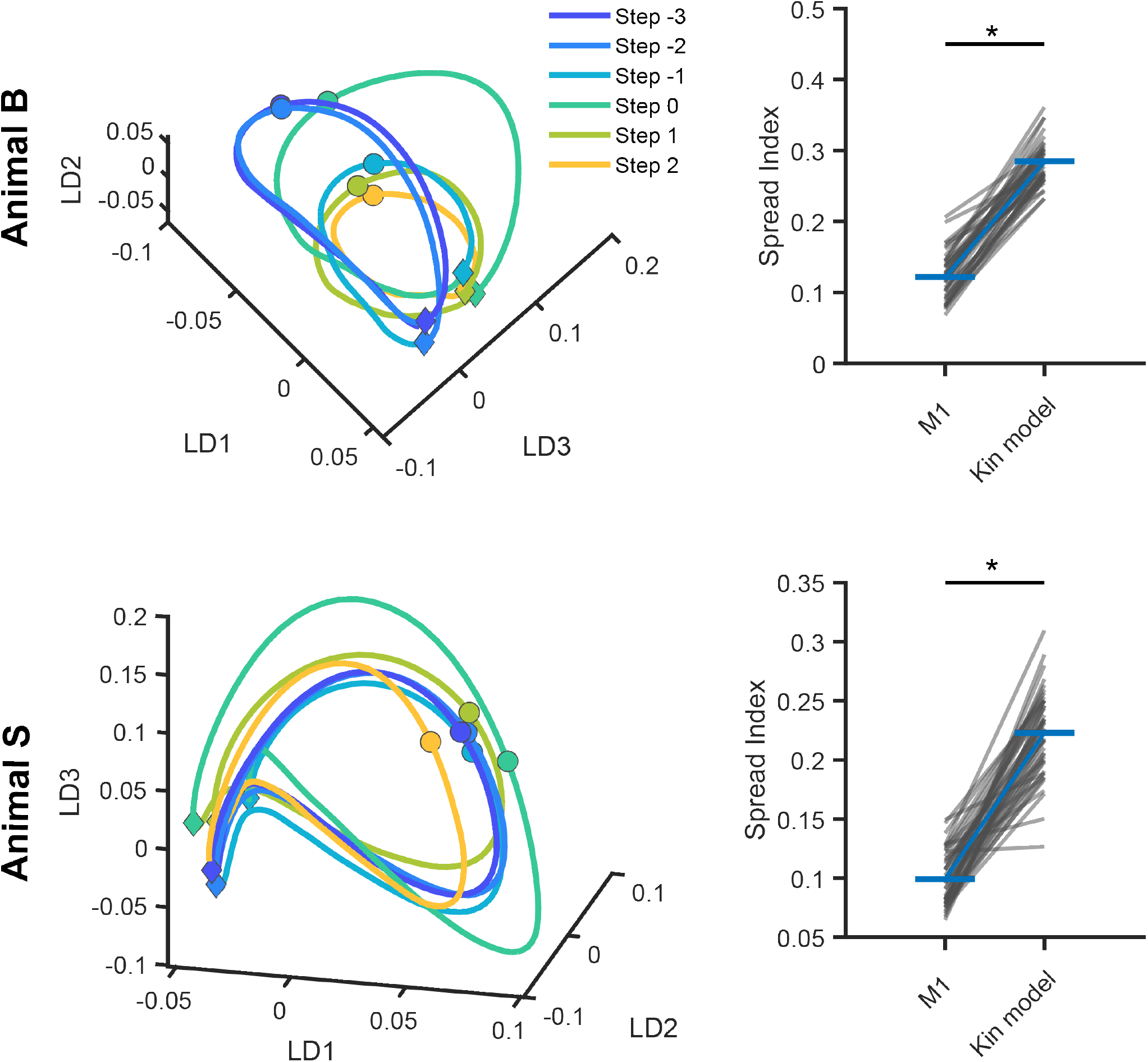
PLDS on model neural data derived from kinematics. For both animal B and animal S, we simulated neural activity using the kinematics and applied our PLDS analysis. Left plots shows the neural trajectories in the first three latent dimensions. There is no projection which results in the same level of overlap across all the strides as in the M1 neural trajectories. Right plots shows the minimum attainable spread metric for either the real neural M1 data, or the kinematic-modeled data.

Projecting the PLDS trajectories along a different angle reveals the modulation of the latent activity in response to the voluntary intervention (figure 7b,f) and the significantly greater spread index compared to the stride-invariant projection (textitp = 4.03×10^−8^ for animal B, textitp = 6.23×10^−9^ for animal S). Like in dPCA, the trajectories appears to diverge away from the basic walking activity (purple and dark blue traces) during the obstacle stride and the preceding stride (light blue and teal traces), before returning to the pre-obstacle region (green and orange traces). These results indicate that M1 maintains a consistent set of rotating neural modes throughout obstacle avoidance stepping while employing a separate set of modes to encode the actual gait adjustment movements. The time evolution of the neural activity in the PLDS latent states as the animal steps over the obstacle is illustrated in movie 2.

### 7.6 Decoding of movement kinematics does not generalize between basic locomotion and obstacle stepping

The existence of neural modes correlating with the efferent intervention of M1 onto the locomotor movements suggests that any kinematic decoders trained on neural data during basic locomotion (and therefore not requiring any gait intervention) would not capture the information represented in these modes. Therefore, we hypothesized that decoders would not be able to generalize across strides during obstacle stepping. To test this hypothesis, we employed a Wiener filter decoder to predict the toe height, the most pertinent kinematic variable for clearing the obstacle, from the neural activity. We trained the decoders with neural and kinematic data from only stride -3 (walk trained) or only stride 0 (obstacle trained) and measured the decoding performance throughout all the strides. We found that while the walk-trained decoder was able to reconstruct the kinematics fairly accurately for the strides before and after the obstacle stride, there was a large amount of error when decoding the swing phase of the obstacle stride (figure 9a,b, blue). Similarly, the obstacle trained decoder was able to accurately decode the toe height during the obstacle stride, but was unable to do so with the surrounding strides (purple).

**Figure 9:**
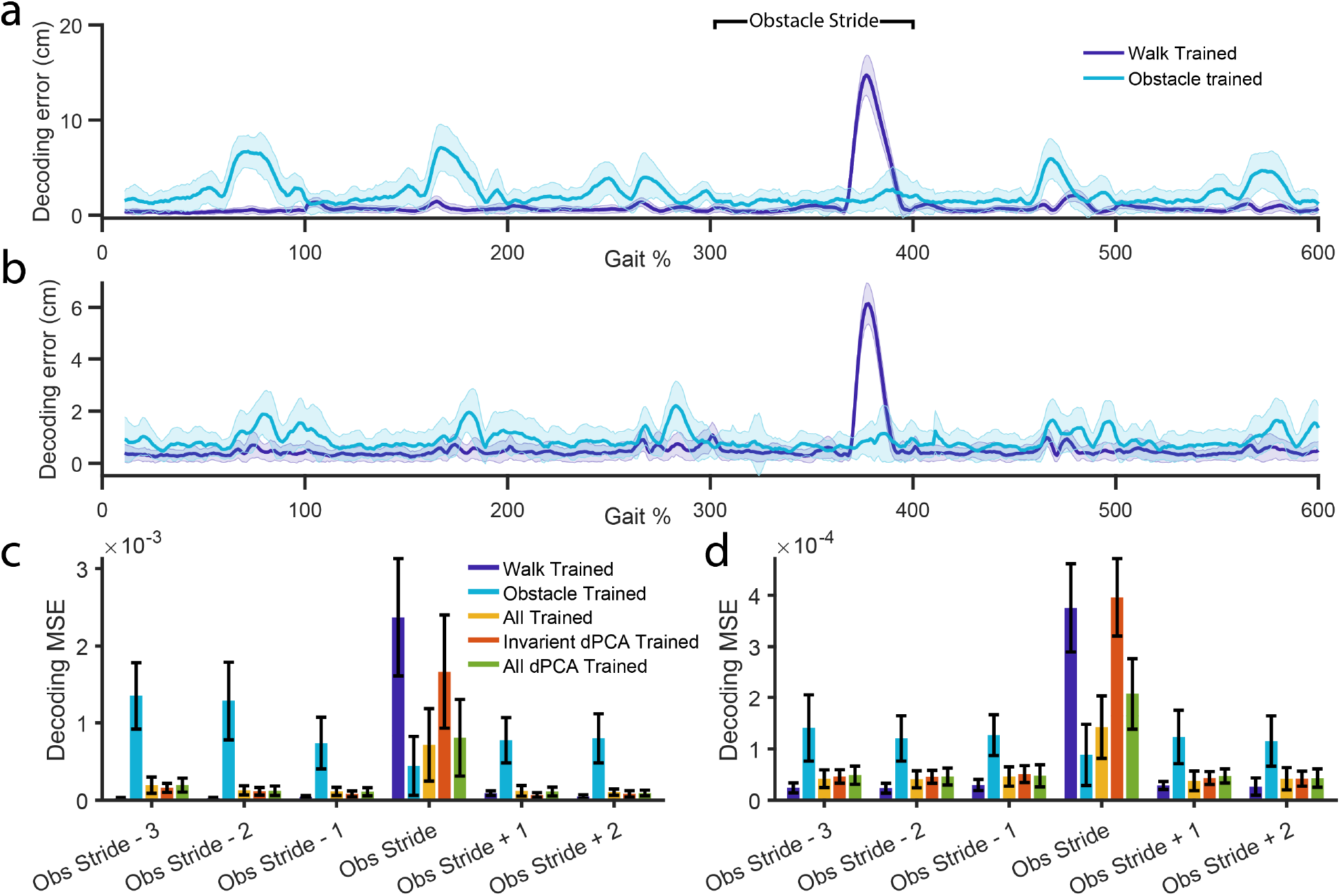
Decoding does not generalize across basic walking and obstacle stepping. **a**,**b)** Error in decoding of the toe height for decoders trained only on the data from the stride before any obstacle movement (Stride -3, shown in purple) or decoders trained only on data from the stride stepping over the obstacle (Stride 0, shown in blue). 300-400% represents the stride over the obstacle, error bars indicate standard deviation. **c**,**d)** Mean squared error for decoding toe height in each of the strides surrounding the obstacle stride, trained on different sets of training data. *Walk Trained* and *Obstacle Trained* are what is shown in **a** and **b**. *All Trained* indicates a decoder trained on data from all of the strides (tested under cross validation). *Invariant dPCA Trained* indicates a decoder trained on data from all of the strides, but only using the stride invariant dPCA components. *All dPCA Trained* indicates a decoder trained on data from all of the strides, and using both the stride invariant and stride dependent dPCA components. **a, c**: animal B, **b**,**d**: animal S.

Finally, to directly test the necessity of the stride-dependent neural modes for capturing the change in kinematics during the obstacle stride, we trained a decoder on neural modes from just the stride-invariant subspace of dPCA, as well as a decoder trained on all neural modes, including those from the stride-dependent subspace. We trained the decoder using the activity from all the strides and performed leave-one-out cross validation. We found that while the decoder trained on just the stride-invariant neural modes performed well on the strides before and after the obstacle stride, it performed poorly on the obstacle stride itself, despite having been trained on data during the obstacle stride. When we included the stride-dependent modes, there was no effect on the first stride (p = 0.1108 for animal B, p = 0.3878 for animal S, Wilxocon rank sum test), but increased the decoding performance during the obstacle stride (figure 9c,d, p <1e-5, wilcoxon rank sum test), suggesting that much of the movement information involved in the gait adjustment is contained in these neural modes.

## 8 Discussion

In this study, we investigated the neural dynamics of primate primary motor cortex during simultaneous execution of two movement modalities - basic locomotion and visually-guided volitional movements in response to an obstacle. Prior research has shown that by segregating the activity of latent neural modes into distinct subspaces, the nervous system is able to carry out multiple neural processes independently within the same population of cells. One previous study has found different M1 subspaces between rodents walking on a treadmill and rodents pressing a lever, and concluded that cortex employs separate dynamics when alternating between these two behaviors(Miri et al., 2017). However, it is still unknown how cortex engages these two different aspects of movement when they have to be carried out simultaneously as a single action. Here, we found that the subspace-partitioning strategy is employed by motor cortex to differentially represent volitional movements and locomotion, behaviors which have been shown to involve different neural processes and structures, when both types movements are being executed at the same time.

In felines and canines, lesion studies have demonstrated that motor cortex is not necessary for the generation of walking movements (G raham Brown, 1911; Grillner et al., 1997), and the control of basic unobstructed locomotion is thought to be managed by subcortical and spinal circuits (Gerasimenko et al., 2006; McCrea & Rybak, 2008). However, volitional movements, such as reaching, do require cortical input. This extends to complex locomotion, as inactivation and lesion studies have shown that without M1 input, animals are unable to precisely place their limbs on a ladder (Beloozerova & Sirota, 1998; Grillner et al., 1997). While it has not been as clearly established in non-human primates that motor cortex merely plays a peripheral role during basic locomotion, numerous studies have suggested that volitional movements and basic locomotion require different amounts of cortical engagement. In prior studies, macaques were able to walk within days after a lesion to the corticospinal tract, but their ability to carry out fine foot movements were almost completely abolished (Courtine, 2005). Similarly, Kuypers and colleagues demonstrated that after a pyramidotomy, macaques could still walk and climb up cages, but lost the ability to make fine dexterous movements (Lemon et al., 2012). These findings suggest that even in non-human primates, regions outside of motor cortex are responsible for, at least partially, generating the movements necessary for locomotion. This is in stark contrast to the essential role that M1 plays during stationary reaching movements. Therefore, the motor cortex of animals carrying out obstacle avoidance must be able to generate the necessary volitional adaptive movements, while simultaneously taking into account the additional movements being generated by external, spinal circuits.

Anatomically, this differential involvement of motor cortex may be instantiated through distinct sub-populations of cells which are activated only during one type of movement. One set of neurons, for example may be dedicated to monitoring the activity of spinal CPGs, while a separate set of neurons direct the timing and magnitude of the movement over the obstacle. However, we did not find evidence for this in our data; the same neural population was active during both locomotion and volitional gait adjustments. This population discharged rhythmically during basic locomotion, but also exhibited an increase in firing activity immediately preceding the increased flexion during the obstacle stride. Additionally, within each behavior, the same recorded population was able decode the limb kinematics during both the basic walking strides as well as the obstacle avoidance strides. Taken together, these results indicate that the same population of cells are generating voluntary movements while also encoding the locomotor rhythm, despite the differential involvement of M1 in these behaviors. However, our findings indicate that the representation of these two movement modalities are segregated within different subspaces within the population.

We employed statistical techniques such as dPCA and PLDS to infer the low-dimensional dynamics in M1 during obstacle avoidance. We note that the state dynamics in the state-space models we employed were linear and may be just an approximation of the true underlying dynamics (Gallego et al., 2017). We then asked - when the animal must carry out a larger movement during the obstacle stride, does the brain modify the whole ongoing neural activity coming from the previous walking strides to carry out the avoidance movement (e.g. figure 1c)? Our results suggests that this is not the case. Both analyses revealed rotational neural subspaces which are unaffected by the change in movement during obstacle stepping, as well as subspaces which were modulated by the gait adjustment and also necessary for accurate decoding of endpoint kinematics during the obstacle step. Therefore, it appears that motor cortex employs a subspace that consistently maintains the same cyclic activity throughout, despite large changes in the movement itself, such as a 2-5x increase in the step height. All the variance corresponding to this large change in movement is confined to its own distinct subspace. This subspace separation is not simply the result of the change in kinematics of the movement, because our control dPCA analysis on the kinematics themselves revealed very small principle angles.

These different subspaces may correspond to different neural processes involved during obstacle avoidance. One strategy that could be employed by the nervous system is to send a feed-forward copy of the motor commands generated by the spinal circuits to cortex (e.g. figure 1a purple arrow), allowing it to generate the necessary volitional movements within the proper context of the ongoing locomotor kinetics. Indeed, it has been previously proposed that the cyclic activity observed in motor cortex during basic locomotion is a reflection of the spinal CPG activity, conveyed through the spinocerebellar tract to the dentate nucleus, and passing through the ventrolateral thalamus before arriving at motor cortex (Beloozerova & Marlinski, 2020; Beloozerova & Sirota, 1998; Marlinski et al., 2012; Massion & Rispal-Padel, 1972). Miri and colleagues have found that this rhythmic activity in cortex during locomotion lies in the null space of volitional movements such as lever pressing (Miri et al., 2017), and therefore this observed activity may serve an ancillary rather than direct role to movement generation. This ancillary role could potentially be to inform cortex of the state of the limb within the gait cycle, should the need for an adaptive movement arise. After all, the amount of lifting required to clear the obstacle is markedly different depending on if the leg is in contact with the ground or at the apex of the swing phase.

The existence of the stride-invariant neural modes we observed are in agreement with this hypothesis. The activity within these neural modes remain consistent throughout all gait cycles, even when the kinematics become drastically different during the obstacle stride. These neural modes also exhibit strong rotational dynamics, consistent with the idea that they are induced by the rhythmic activity in spinal CPGs. Additionally, these neural modes were sufficient for decoding end-point kinematics during basic locomotion, but failed during the volitional gait adjustment, suggesting that they do not carry information involving the adjusting movement. However, further experiments employing causal circuit perturbations will be needed to truly determine the origin of these signals in non-human primates.

Meanwhile, the variance related to volitional gait-adjusting movement appear to be contained to a separate subspace defined by a separate set of neural modes. The integration of the visual information along with the calculation of the precise movement required to clear the obstacle may be subserved by these modes. Our results do not preclude the possibility that other factors may be present in these stride-specific subspaces, such as attention or postural changes not observed in our kinematic analysis. Additionally, we note that while the neural dynamics of M1 contain both sets of subspaces, M1 may not be solely responsible for carrying out the integration of the volitional movement with the ongoing locomotor movements. Other upstream regions such as premotor cortex (Nakajima et al., 2019), parietal cortex (Marigold et al., 2011; Marigold & Drew, 2011), or ventrolateral thalamus (Beloozerova & Marlinski, 2020) may also contribute to the integration computations before relaying the signals to M1.

Finally, our findings have implications for the development of brain-machine interfaces (BMIs) aiming to restore hindlimb functionality for patient with motor deficits. In recent years, advancements have been made in the development of closed-loop systems for restoring walking ability after spinal cord injury (Capogrosso et al., 2016; Donati et al., 2016). These systems utilize electrophysiology recordings from cortex to drive either spinal stimulation or movement of an exoskeleton, and have demonstrated remarkable success in allowing subjects to recover locomotor ability. These systems currently only aim to restore basic locomotion, and not precise directed leg movements. Our decoding results suggest that future BMIs aiming to restore a wide range of hindlimb movements, including visually adaptive locomotion, should consider including both basic locomotion as well as volitional movements during decoder calibration in order to achieve optimal performance.

## 2 Acknowledgements

This work was sponsored in part by the Defense Advanced Research Projects Agency (DARPA) BTO under the auspices of Dr. Doug Weber and Alfred Emondi through the [Space and Naval Warfare Systems Center, Pacific OR DARPA Contracts Management Office] Grant/Contract No. D15AP00112 (DAB), by the Merit Review Award #I01RX002835 from the United States (U.S.) Department of Veterans Affairs, Rehabilitation Research and Development Service (DAB). The contents of this manuscript do not represent the views of VA or the United States Government. The Department of Veterans Affairs did not provide any direct support of the animal work conducted for this project. We would like to thank Ellen Xing for her artistic contribution in the creation of figure 1 and figure 2. We would also like to acknowledge the Pablo J. Salame ‘88 Goldman Sachs endowed Associate Professorship of Computational Neuroscience at Brown University (WT), the Howard Reisman ‘76 Family Graduate Fellowship Fund, and the Charles A. Dana Graduate Fellowship Fund (DX). These funding sources had no involvement in the design of this study, collection or analysis of data, nor in the authorship of this manuscript.

## 9 Author Contributions

DX and DAB conceived and designed the experiments and performed the implantation surgeries. DX carried out the experiments and performed the data analysis with input from WT and DAB. DX wrote the manuscript with input from WT and DAB. All authors edited the manuscript.

## 10 Data availability statement

The data that support the findings of this study are available from the corresponding author upon reasonable request

## 11 Code availability statement

Peer-reviewed, published libraries for dPCA, jPCA, and circular statistics are referenced in the text at appropriate locations. The analysis code that supports the findings of this study is available from the corresponding authors upon reasonable request.

---

Movie 1: Stick figures illustrating the hindlimb kinematics across all trials for basic unobstructed walking (red) or the step over the obstacle (blue) in animal B. Video illustrates the kinematics before they were normalized to the gait cycle. Trials were time-locked to the start of the swing phase (toe off event).

Movie 2: PLDS neural trajectories during obstacle avoidance. This movie illustrates the time-varying neural activity in the first three PLDS latent dimensions (LDs) of animal B. The obstacle stride, three strides before the obstacle stride, and two strides after the obstacle strides are shown. Thin traces represents the activity for each individual trial. Thick traces represents the trial-averaged activity, e.g. what is displayed in figure 5a-b.

